# CloudBrain: Online neural computation in the cloud

**DOI:** 10.1101/2021.01.21.427662

**Authors:** Leon Bonde Larsen, Rasmus Karnøe Stagsted, Beck Strohmer, Anders Lyhne Christensen

## Abstract

Neuromorphic computing currently relies heavily on complicated hardware design to implement asynchronous, parallel and very large-scale brain simulations. This dependency slows down the migration of biological insights into technology. It typically takes several years from idea to finished hardware and once developed the hardware is not broadly available to the community. In this contribution, we present the CloudBrain research platform, an alternative based on modern cloud computing and event stream processing technology. Typical neuromorphic design goals, such as small form factor and low power consumption, are traded for 1) no constraints on the model elements, 2) access to all events and parameters during and after the simulation, 3) online reconfiguration of the network, and 4) real-time simulation. We explain principles for how neuron, synapse and network models can be implemented and we demonstrate that our implementation can be used to control a physical robot in real-time. CloudBrain is open source and can run on commodity hardware or in the cloud, thus providing the community a new platform with a different set of features supporting research into, for example, neuron models, structural plasticity and three-factor learning.

## 1 Introduction

In traditional artificial neural networks (ANNs), the activation of neurons is represented as a scalar and information is propagated through the network in discrete steps. While this model can be computed efficiently on standard CPUs and GPUs, it is a very simplistic abstraction of biological neural networks. Biological neurons primarily communicate asynchronously through action potentials or *spikes* (Sterling and Laughlin, 2015). The timing of spikes can encode crucial information, for example in the auditory system where temporal information is used to infer direction of a sound source (Carr and Konishi, 1990, Haessig et al., 2020). Spiking neural networks (SNN) are a class of ANNs in which the temporal aspects of interneuron communication are explicitly considered: neurons asynchronously produce and communicate via discrete events (spikes), and SNNs thus allow encoding of information in the timing of the events.

Different spiking models have been described in literature ranging from the relatively simple integrate-and-fire model (Keat et al., 2001, Jolivet et al., 2004, Paninski et al., 2004) to the more complex Hodgkin-Huxley model (Hodgkin and Huxley, 1952). To improve biological fidelity there is, however, still a need for experimenting with new models. For example biological neurons can be non-spiking (Sterling and Laughlin, 2015) and there could be advantages of combining spiking and non-spiking models (Woźniak et al., 2020). Integrating non-spiking neurons in SNNs can be a biologically plausible way to interface analogue sensors and can provide more control over network behaviour (Strohmer et al., 2020).

Current implementations of learning, both in ANNs and SNNs, are based almost exclusively on adapting parameters in the synapses connecting the neurons. Such synaptic learning also plays a crucial role in biological learning, but in ensemble with for example structural adaptations of the network and influence from different neuromodulators providing reward signals or adapting neuron behaviour (Sterling and Laughlin, 2015, Price et al., 2017). In a neuromorphic engineering context, structural plasticity has been shown to improve facilitation of the hardware (Qi et al., 2018), increase success of learning a task in reinforcement learning (Spüler et al., 2015), and in unsupervised classification tasks (Roy and Basu, 2017). Online adaptation of network structure is, however, not directly supported in current neuromorphic hardware, thus restricting research to pure simulations.

In this paper, we present the CloudBrain platform for simulating SNNs. CloudBrain utilises modern cloud technology to create an infrastructure capable of executing SNNs in a computer cluster. This gives several advantages: 1) There are practically no constraints on the model elements. If the concept can be described in code it will also run in the cluster. 2) It allows access to all events and parameters both online and offline, making it easier to monitor, develop and test solutions. 3) The structure of the network can be reconfigured online allowing model elements to affect connectivity. 4) The network can run online and control a robot through the *robot as a service* principle (Kuffner, 2010) to interact with the environment. 5) It runs on standard computers and is based on well-documented, field-tested, free, open source software. We present the architecture of CloudBrain, demonstrate the advantages of the approach, and deploy it in closed-loop control of a robot.

### 1.1 SNN simulators

SNNs can be simulated on PCs or supercomputers using specialised software such as the GENESYS (Bower et al., 2003) and NEURON (Carnevale and Hines, 2006) simulators or the more computationally tractable NEST (Gewaltig and Diesmann, 2007), BRIAN (Stimberg et al., 2019) and CARLsim (Chou et al., 2018). These simulators are very useful for investigating the behaviour of networks and of their constituent parts. Their limitation lies in not being able to embody the neural simulation for instance to control a physical robot. It has also been suggested that they are less suited for evolving models (Nowke et al., 2018) because the model requires external control while evolving.

An alternative to the software simulations is neuromorphic hardware. Since a spiking neuron only needs to do work whenever it receives a spike, it can operate asynchronously. That observation has been the basis for developing non-von-Neumann computer chips (Furber, 2016) leading to small, scalable, fast and energy-efficient devices for researching and deploying SNNs, such as SpiNNaker (Furber et al., 2014) and BrainScales (Schemmel et al., 2010) developed in the Human Brain Project, IBM’s True North (Essera et al., 2016), Loihi from Intel (Davies et al., 2018), and the analog DYNAPs (Qiao et al., 2015, Moradi et al., 2018) developed at ETH Zurich. Such neuromorphic hardware is well suited for embodied experiments since the neuronal computations can run in real time, for example controlling a physical robot.

Each chip represents a trade-off between features and limitations. Common for all of them is that the interface to and from the chip is a bottleneck and does not allow the user to export all the spike events happening in the chip. This can complicate monitoring and makes it harder to analyse a network. The SpiNNaker platform (Furber et al., 2014) is available for loan but otherwise gaining access to neuromorphic hardware can be challenging because only few units exist or because intellectual property rights restrict its use. Support in the form of software frameworks and documentation can also be limited as is support for special features, for example to investigate new neuron models.

More flexible hardware implementations of SNNs have been demonstrated on Field Programmable Gate Arrays (FPGAs) and Graphics Processing Units (GPUs). FPGAs are programmable devices consisting of numerous logic blocks that can be almost arbitrarily connected. Once programmed, the FPGA’s performance is comparable to specialised chips. The use of FPGAs to simulate SNNs has been found to be highly scalable (Moore et al., 2012, Wang and van Schaik, 2018) and some work suggests vector processing implemented in FPGAs can help mitigate the memory bottleneck problem that reduces access to spikes and parameters (Naylor et al., 2013). However, the lack of hardware support for floating-point arithmetic limits FPGAs to the simpler neuron models and they do not easily support online changes to network structure.

FPGAs are still quite uncommon and programming them is very different from computer programming so their use requires special training. GPUs, on the other hand, are common in most modern PCs and implementations of SNNs on GPUs have been demonstrated to be highly scalable (Hoang et al., 2013, Chou et al., 2018). GeNN, a GPU-enhanced simulation software based on NVIDIA CUDA technology, even out-performed some state-of-the-art specialised chips with regards to speed and power-consumption (Knight and Nowotny, 2018). The availability of embedded GPU platforms, such as NVIDIA’s Jetson TX2 also enables GeNN to be used interactively to control a robot. The strong commercial development of GPUs is constantly moving the boundaries for what is possible but generally, moving data to and from the GPU memory is a bottleneck limiting access to spikes and parameters.

Sometimes hybrid systems can enable new features or remove limitations. For example the SpiNNaker million-core machine is available through the Human Brain Project’s portal (Human Brain Project, 2017) for running even very large simulations and Intel Labs developed a cloud-based platform for research community access to scalable Loihi-based infrastructure (Intel, 2019). However, none of them support online experimentation, for example with robots. Brian2GeNN (Stimberg et al., 2020) is a software package that uses GeNN to accelerate simulations defined in Brian on GPU hardware. GPU acceleration of simulation software has also been demonstrated to improve performance on supercomputers and enable larger simulations on single computers (Hoang et al., 2013, Chou et al., 2018).

### 1.2 Cloud infrastructure

In recent years, cloud-based technology has seen rapid development driven by the demand for distributed and highly scalable IT-solutions (**?**). When executed in the cloud, a computer program often runs in a virtual environment called a *container*. Seen from the program the container is like any computer with resources such as CPU, memory, disk and an operating system while in fact the containers share these resources. Asynchronous, event-based architectures in particular have excelled in order to handle millions of social media users or e-commerce transactions. Programs are often asynchronous, meaning that they are waiting for input for example from a user requesting a website or completing a purchase. While waiting the program needs no computing resources and thus other programs in other containers can use the hardware. This fits well with the SNN model, where the neuron only does work when an input event is present.

Modern IT solutions generate a lot of data and it is common to handle it as event-streams (**?**). Event-streams are ordered in topics such that nodes subscribing to a topic receive events that are published on that topic. There can be many publishers and many subscribers to a topic and there can be many topics. Copying and distribution of the events are handled by highly optimised and extremely scalable infrastructure software making it easy to interface programs with the event-stream. Such an architecture is well suited for handling spike events since many neurons need to receive the same events.

## 2 Architecture

An SNN simulation in CloudBrain is essentially a collection of small programs implementing mathematical models and communicating the resulting events as they happen. CloudBrain is the platform that runs the programs in a scalable and modular way. In CloudBrain, both neurons, synapses and any other models are user-defined programs, while spikes, parameters, and any other messages are events consisting of a timestamp and an arbitrary payload.

### 2.1 Programs

The *NeuronProgram* and *SynapsePrograms* run under the *ControlProgram* in order to hide the complexity of communication and OS-specific interfaces. One *ControlProgram* can run one *NeuronProgram* and multiple *SynapsePrograms* and to ensure scalability, the *ControlProgram* is executed in a container. Thus the *ControlProgram* can run on any host within the cluster and multiple *ControlPrograms* can run on the same host (figure 1). The user controls if the *ControlProgram* runs synchronously updating the neuron model at a specific rate or asynchronously only updating the model when a spike is received. The same goes for the *SynapseProgram* which is responsible for keeping information about the connections (for example weight and delay) and attaching it to the payload of received spikes before handing them to the *NeuronProgram*. The connection information is provided when the connection is first made but can also be updated during execution. *Events* are implemented as asynchronous messages transmitted following the publish-subscribe pattern (Birman and Joseph, 1987) so a *ControlProgram* receives messages only from topics it has subscribed to. Topics contain either *ControlEvents*, handled by the *ControlProgram* or *NeuroEvents*, passed first through a *SynapseProgram* and then handled by a *NeuronProgram*. Each *ControlProgram* has an individual *ControlTopic* that it always subscribes to while all other subscriptions are set up at run-time.

**Figure 1:**
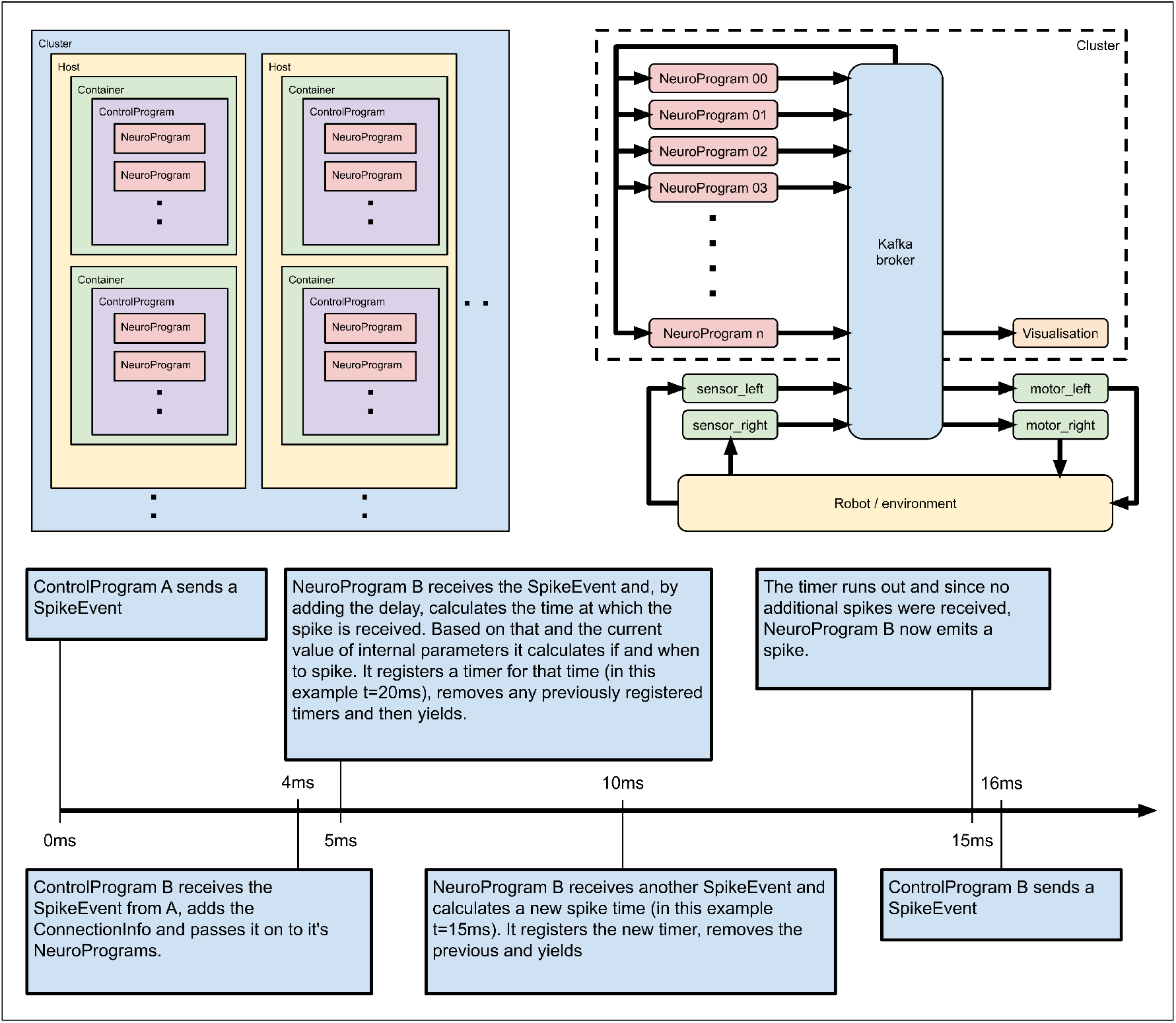
**Top left**: Shows how CloudBrain scales. Several *NeuroPrograms* can run under one *ControlProgram*. Several containers, each running one *ControlProgram*, can run on one host and the cluster is made up of any number of hosts. **Top right**: Conceptual overview of the robot experiment. The *NeuroPrograms* communicate through the kafka broker with the sensor and motor populations on the robot. **Bottom**: Simplified scenario demonstrating how asynchronous neuron models are implemented in CloudBrain.

### 2.2 Global control

A *GlobalController* is responsible for scaling the number of containers in the cluster and for configuring the *ControlPrograms* by emitting *ControlEvents*. Each *ControlProgram* in the cluster has a unique ID known to itself and the *GlobalController*. To set up an experiment, the user writes a *GlobalController* program that tells each of the *ControlPrograms* which *NeuronProgram* and *SynapsePrograms* to run, which parameters to use and how to connect. It also defines any groupings of *NeuronPrograms*, for example into populations. If non-standard neuron models are used, they have to be implemented in code and either provisioned to the cluster before running the *GlobalController* or sent to the individual container using control events, during execution. The *GlobalController* can be run from any computer on the same network as the cluster.

### 2.3 Simulation method

Ideally, the *NeuronPrograms* and *SynapsePrograms* should be asynchronous, meaning that they only use processing resources when they receive or transmit an *Event*. This greatly improves performance but is not practical for all neuron models and thus they can also be periodic. To run asynchronously, every time a neuron receives a spike, it must predict when it will spike based on the mathematical model of the neuron and its internal parameters. The neuron then registers a timer to wake it at that time and removes any previously registered timers (figure 1).

Communication in the cluster is faster than in biological neurons. Because the *NeuronPrograms* calculate behaviour based on timestamps and not on the actual time of arrival, the cluster essentially spends the time accounted for in the biological delays to complete the required calculations and communicate the results. If that time is insufficient, the post-synaptic neuron will know that a deadline was missed and can report it. The *SynapseProgram* receives incoming *SpikeEvents* and attaches the connection information (typically weight and delay) before passing them on to the *NeuronProgram*. When the *NeuronProgram* emits a spike, it also alerts the *SynapseProgram* thus allowing it to update the connection parameters according to its learning rule.

## 3 Methods

A proof-of-concept implementation based on Python and open-source software was built in order to validate the proposed architecture. The code is available on a git repository along with a demo of CloudBrain running on a single PC and instructions on how to set up a cluster with a cloud provider. All is available at sdu.dk/cloudbrain.

### 3.1 Implementation

*ControlProgram* and *NeuronProgram* are written in Python. A custom *NeuronProgram* inherits from a base class so the user only needs to override the functions used and need not care about the inner workings of the *ControlProgram*. The functionality exposed by the *NeuronProgram* base class include methods to emit events and to register callback functions for event reception or the expiration of timers. The Python code is integrated in a minimalistic docker image (**?**) and executed in docker swarm (**?**). Using the continuous integration tools included in gitlab (**?**), the process of deploying the code on the docker swarm is automated. Log activity from the containers is collected using Filebeat (**?**), allowing the logs to be searched and viewed from a web-based interface. Events are handled by Apache Kafka (**?**), an open-source stream-processing software platform, providing high-throughput and low-latency communication. Messages are JSON encoded and consist of a timestamp, sender ID and an arbitrary payload. Since Kafka is agnostic to the payload, it can be seamlessly changed to fit any computational model, to set parameters in a *NeuronProgram* or to retrieve arbitrary information. One of the great advantages of an event-based system is the availability of tools to view, search and aggregate data. We use Elasticsearch and the visualisation dash-board Kibana. This allows for fast, online and virtually unconstrained visualisation of activity in the network, for example, to monitor spiking rates at population level or for individual neurons, to analyse behavioural patterns or to monitor the flow of *ControlEvents*.

### 3.2 Hardware

The on-premises cluster consists of 15 PCs, each with 2 Intel Xeon 2.55GHz cores, 8 GB RAM, Gigabit network and with Debian9 installed on a solid state drive (figure 5 shows a photo of the cluster). An additional PC with Intel Xeon 2.67GHz quad-core, 12 GB RAM and SSD is used to run Kibana, Kafka, Elasticsearch connector and Zookeeper (used by kafka). Another PC with Intel I5 3.30 GHz quad-core, 12 GB RAM and SSD is used to run Elasticsearch. Elasticsearch runs on a separate host to keep peak CPU and memory usage from interfering with the performance of Kafka. Finally a PC with Intel I5 2.90 GHz quad-core, 16 GB RAM and SSD was used to monitor the cluster using Grafana and InfluxDB. On each of the hosts in the cluster, Telegraf was installed to collect information about the utilisation of the nodes. The machines are connected to a gigabit managed switch using Cat5e cables and communication to the robot is provided by a VPN tunnel through a Wi-Fi access point. As an alternative to procuring an on-premises cluster, we repeated the experiments with Google Cloud Platform (GCP) executing CloudBrain. 14 virtual machines were configured each with 4 vCPUs running at 2 GHz, 15 GB RAM and 100GB disk. 10 VMs were used to run the neurons and the rest were used as VPN gateway, Kafka broker, Elasticsearch and connectors to Elasticserch. All VMs were located in Finland while the robot was in Denmark.

### 3.3 Robot platform

The mobile robot described in Larsen et al. (2013) is optimised for rapid prototyping and consists of a wooden board with two rear wheels connected to motors and a castor wheel in front. In this work it was fitted with two custom bumper sensors, one on each side of the front (figure 5). An on-board RaspberryPi 3 model B connected to the cluster via Wi-Fi is responsible for generating PWM signals for the motors based on received events and for emitting events based on the state of the sensors. A piece of software running in the cloud translates spikes into motor messages and sensor messages into spikes. An H-bridge supplies the current to the motors based on the generated PWM signals. The two sensors are implemented with micro-switches and each sensor emits spikes on its own topic. When the sensor is activated, it emits spikes with a frequency of 100Hz and when the sensor is not activated it emits spikes with a frequency of 10Hz. Each motor has its own topic and the output is controlled by a running average of the number of spikes received within the past 50ms. The average is then linearly mapped from 1-4 spikes to ± 2 wheel rotations per second, controlled by a standard PID controller.

During experiments the robot was enclosed in a 2.5m × 1.5m environment with slanted 25cm corners (see centre insert of figure 2). The environment can be configured as *corridor* or *box* by adding or removing the rectangle in the centre. The experiments were repeated several times in each configuration. The controller for the robot is based on the Braitenberg principle (Dennett and Braitenberg, 1986) where the left sensor has an inhibitory effect on the right motor and vice-versa. The network consists of six populations, each with five neurons (figure 2). Apart from the two sensor populations (A and B) and the two motor populations (C and D), an excitatory bias (E) is provided to both motor populations to make the robot move, and the last population (F) injects noise into the system to make the neurons fire out of phase.

**Figure 2:**
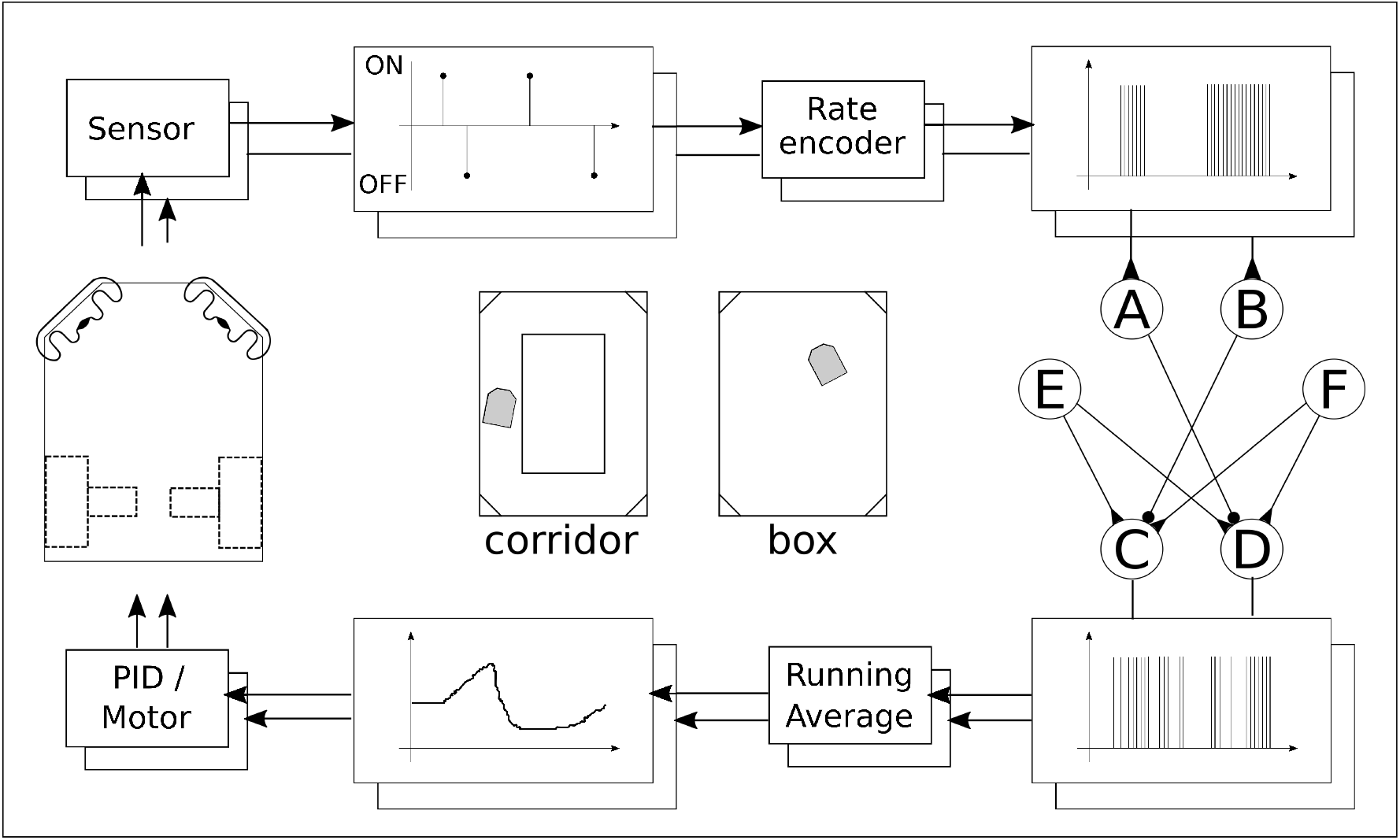
The control loop of the robot. Onset/offset events from the sensors are rate encoded and fed into populations A and B, respectively. E is injecting a bias (to keep the robot moving) and F injects noise. The motor neuron populations C and D are decoded by a running average and used as set-point in the PID for the motors. The centre insert shows the corridor track and the box track used for testing the robot.

### 3.4 Neuronal models

We implemented and tested three popular neuron models: Integrate-and-Fire (IF) (Keat et al., 2001, Jolivet et al., 2004, Paninski et al., 2004), Leaky-Integrate-and-Fire (LIF) (Stein, 1967, Tuckwell, 1989) and Adaptive-Exponential-Integrate-and-Fire (AEIF) (Brette and Gerstner, 2005, Gerstner and Brette, 2009). The IF and LIF neurons are implemented fully asynchronously so the neurons only use computational power when input spikes arrive or the neuron emits a spike. The AEIF model consists of two coupled differential equations that need to be integrated over time. To keep it asynchronous would make it computationally heavy because it would need to recalculate next spike time and set the timer accordingly every time a new spike arrives. Instead the model was implemented synchronously using the looping function with an update frequency of 1ms. We implemented the Spike-Timing-Dependent Plasticity (STDP) rule, such that it runs asynchronously and updates the weight of the synapse.

## 4 Results

To support the claims that CloudBrain provides 1) no constraints on the model elements, 2) online reconfiguration of the network, 3) online operation, and 4) access to all information, we provide three demonstrations. We demonstrate 1) the implementations of three popular neuron models and a synapse model, 2) that the morphology of the network can be changed during operation, and 3) that a robot can be controlled online by CloudBrain running on a cluster. Furthermore, to demonstrate how all information in the SNN can be monitored online, we provide the live plots from Kibana. Finally, we evaluate timing and load both on our own cluster and with a cloud provider.

### 4.1 Model implementations

To demonstrate the implemented neuron models, two different experiments were made. In the first one, a spike source was connected to the IF and LIF, respectively and their voltage potential and firing was observed (figure 3, top). The voltage potential of the IF neuron increases linearly until it fires, whereas the voltage potential of the LIF neuron charges exponentially. The AEIF model was tested in the regular bursting mode with constant current input (Naud et al., 2008) using parameters from NEST (Gewaltig and Diesmann, 2007) while voltage potential, adaptation variable and spiking pattern were observed (figure 3, middle). The voltage potential of the AEIF neuron depends on the adaptation variable and, as it falls to a certain level, the neuron bursts. This confirms normal behaviour for all three neuron models.

**Figure 3:**
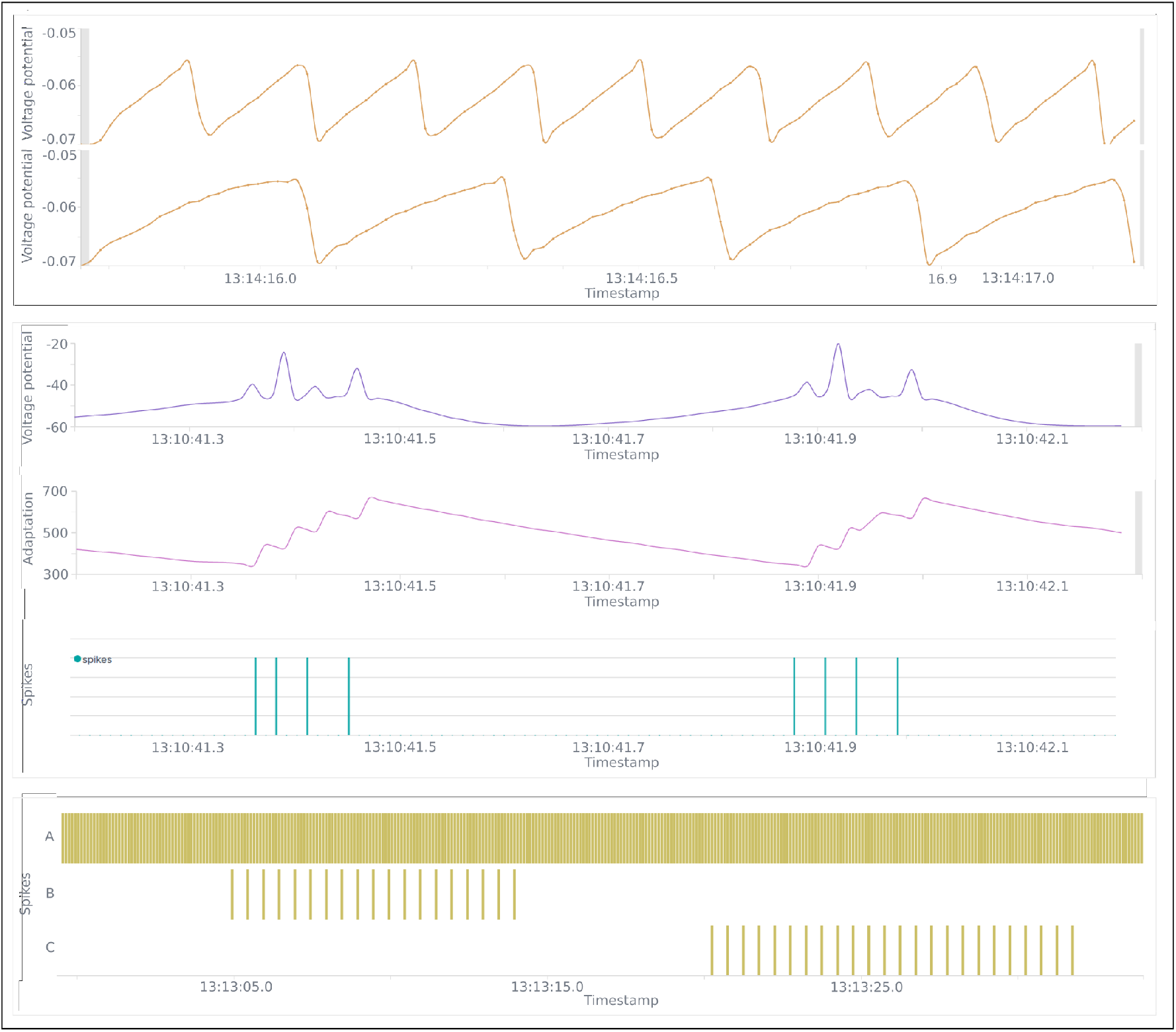
Screenshots from live monitoring in Kibana. **Top**: Voltage potential from experiment with Integrate-and-fire and Leaky-integrate-and-fire. **Middle**: Voltage potential, adaptation variable and spikes from experiment with Adaptive-exponential-integrate-and-fire. **Bottom**: Spikes from morphology experiment showing spikes from the spike source (top line) and from the two Integrate-and-fire neurons. The spike source is in turn connected to and disconnected from the neurons, causing them to spike.

Synaptic learning is demonstrated in a setup where a noisy spike source has excitatory connections to two LIF neurons. The two LIF neurons are connected with a synapse using STDP on all pre- and post synaptic events. The noisy spike source emits events with a random time difference between 100 ms and 1 s. Every time one of the LIF neurons spikes, the STDP synapse will update its weight based on the time since the other neuron last spiked. Figure 4 shows the evolution of synaptic weight in an STDP synapse. From the zoomed-in figure, we see that when a pre-synaptic spike occurs (green vertical line) the weight increases by an amount inversely proportional to the time since last post-synaptic spike. Similarly at the post-synaptic spike times the weight is reduced inversely proportional to the time since last pre-synaptic spike.

**Figure 4:**
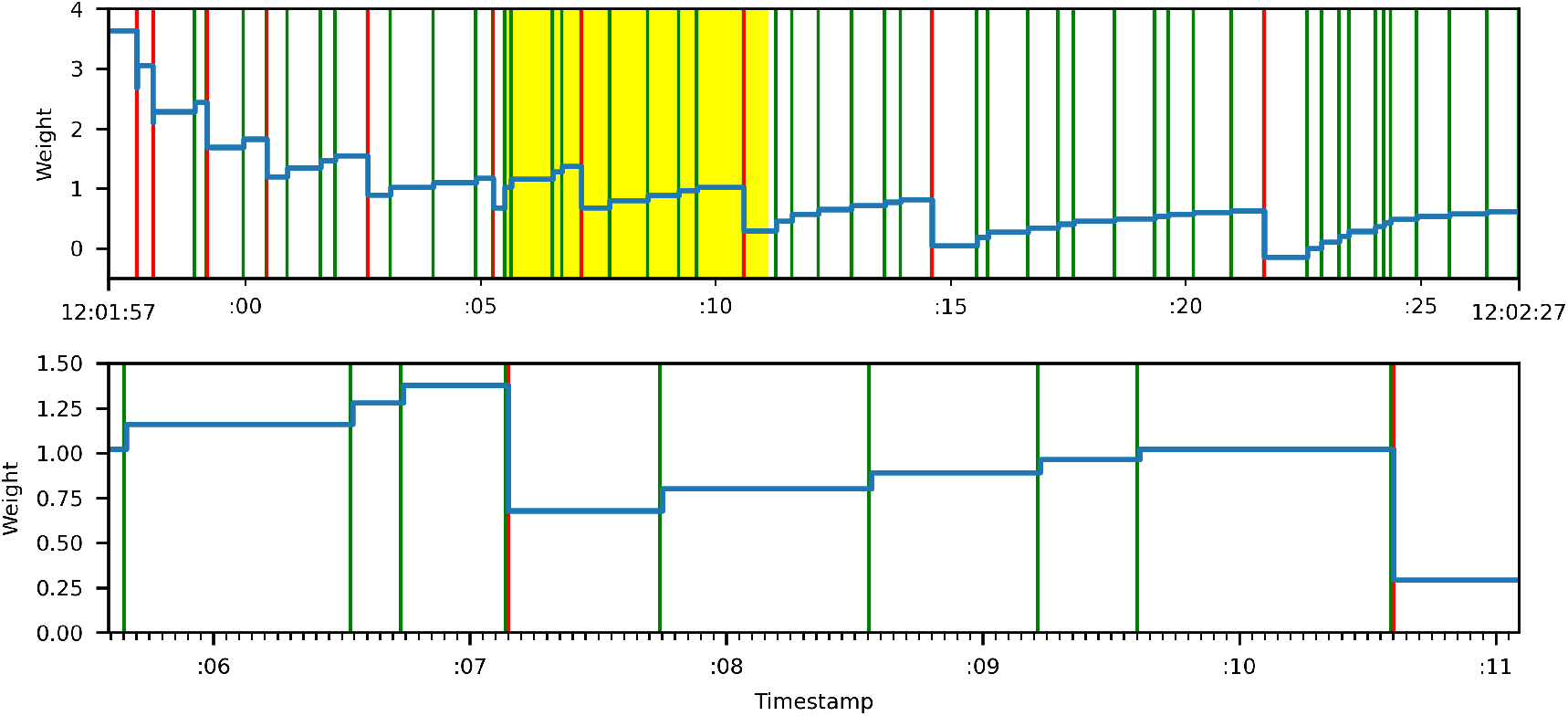
Synapse learning using STDP. Green vertical bars represent pre-synaptic spikes, while red vertical bars represent post-synaptic spikes. The blue line represents the synaptic weight. The bottom plot is a magnification of the yellow area in the top plot.

### 4.2 Morphology

To demonstrate that the network can be reconfigured online, an experiment was made with a 10Hz spike source (A) and two IF neurons (B and C). After 5 seconds, the source is connected to neuron B thus making it fire. After another 10 seconds, the connection is removed and the same procedure is repeated for neuron C. We plot the spiking pattern of the three neurons in figure 3. The morphology changes shown here are very basic, but demonstrate that connections can be created, updated or removed by sending ControlEvents.

### 4.3 Online operation

To demonstrate online operation, two experiments with the robot are reported, each in a different environment. The spiking rate of the populations were plotted online in Kibana and the resource use in the cluster were plotted using Grafana. Here we provide screenshots from both and a video of the experiment is available in the supplementary material. The experiments were run on the on-premises cluster and repeated in the GCP cluster.

To evaluate the total delay in the robot experiments, we used videos to analyse reaction times when the robot is colliding with the wall. We also estimated the delay from a change in spike rate of the sensor population until the corresponding change in the opposite motor population. This represents the total delay in the neuronal system without the robot. The delay of a single event through Kafka was measured by subscribing to a topic and publishing 100 events on it. This was measured both over WiFi and within the clusters. We report the median of the 100 measurements along with the 1st and 99th percentiles.

The robot was able to negotiate both corridor and box, reacting quickly and displaying the expected Braitenberg behaviour. The total delay from sensor activation to motor reaction was estimated based on several collisions with the wall, recorded on video. Upon collision the robot was thrown back a few centimetres and the controller reaction was fast enough to prevent the robot from hitting the wall again. It took on average 5 frames (min. 4 frames, max. 8 frames), corresponding to 166 ms (min. 133 ms, max. 266 ms) to react. There are different contributions to this delay, mainly the characteristics of the motor. The total delay from a change in a sensor population to the corresponding change in the motor population could not be determined precisely due to noise but was estimated to 30–80 ms based on the plots from Kibana (figure 5). The delay of a single event through Kafka was measured over 100 messages. The time from an event was sent until it was received was 24 ms via Wi-Fi (median: 24.0 ms, 1st: 8.0 ms, 99th: 320.4 ms) and 4.2 ms within the cluster (median: 4.2 ms, 1st: 2.7 ms, 99th: 5.5 ms).

**Figure 5:**
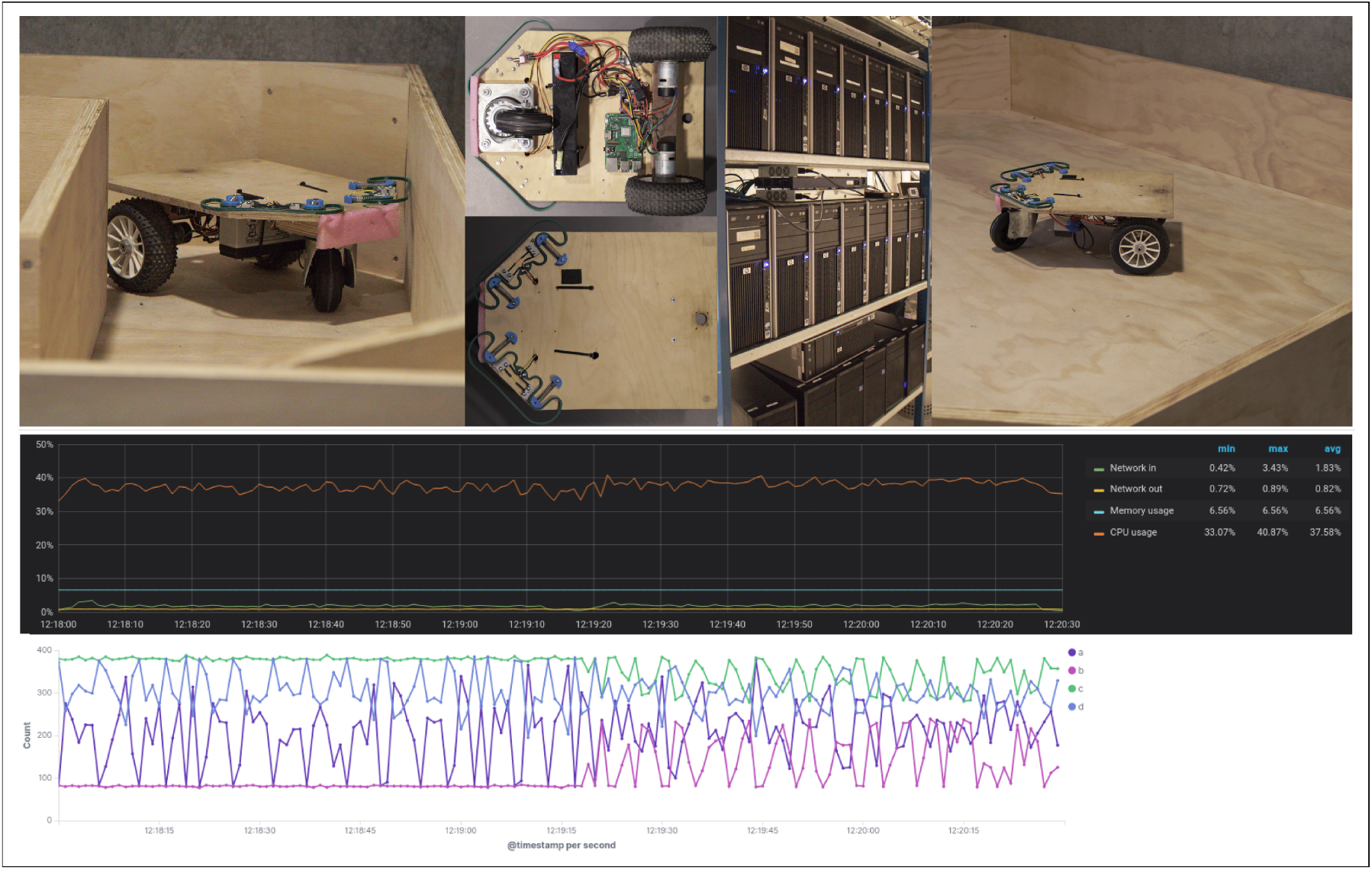
**Left image**: The robot in the corridor. **Right image**: The robot in the box. **Middle images**: The robot seen from below, the robot seen from above and the on-premise cluster. **Top plot**: Load on the cluster during operation. All values are taken as the percentage of the cluster’s total capacity. The plot is taken directly from Grafana as displayed live. **Bottom plot**: Spike rates on the interface populations (a/b for left/right sensor and c/d for left/right motor). The first half is in the box and the second half is in the corridor. While driving in the box, the robot displays a wall following behaviour thus always activating the same sensor and reacting with the same motor. The plot is taken directly from Kibana as displayed live.

Figure 5 also shows the load on the cluster while the robot is navigating the corridor. The memory usage is constant because once the neurons are running they do not make additional allocations. The network usage is less than 2%. For comparison, the idle cluster had memory usage of 6%, CPU usage 3% and network 0%. Under the experiment an average of 1930 spikes per second were passing through Kafka. Benchmarking of the Kafka broker showed that it could handle approximately 400.000 spikes per second.

The same experiment was conducted on GCP for a duration of 15 minutes. Similar to the delay measured with the on-premises cluster, the latency was measured from the robot to the GCP. The delay measured to 47.3 ms (median: 46.7 ms, 1st: 46.2 ms, 99th: 52.6 ms). The spike delay within the cluster was measured to 3.5 ms (median: 3.2 ms, 1st: 2.7 ms, 99th: 5.5 ms).

The *ControlProgram* allows the user to toggle live logging of internal variables and parameters. The values are sent, as events, only when they change. The plots in figure 3 and 5 are all live screenshots from Kibana demonstrating that parameters are available online. Data can also be accessed online without Kibana for more advanced plots or extracted for offline generation of plots better suited for publication.

## 5 Discussion and conclusion

In this contribution, we propose a novel architecture for simulating spiking neural networks on readily available cloud infrastructure. We describe a proof-of-concept implementation, demonstrating how neuron and synapse models can be implemented and how the network connections can be updated while the system is running. Finally we demonstrate how a Braitenberg-controlled differential drive robot could be operated online from both our on-premises cluster and from a Google Cloud Platform cluster.

The numbers provided are highly dependent on hardware, application and other implementation specific factors. We therefore provide the numbers observed in our proof-of-concept implementation without expecting them to generalise perfectly to other implementations. Our on-premises cluster built from 15 refurbished PCs over ten years old was able to support the experiments in this paper and the total cost of running the robot experiment for 15 minutes on GCP was approximately 2 USD. This shows that hardware for this type of simulation is readily available.

We did experience some jitter in the delay times and occasionally an event was late by a factor of hundreds which means it missed the deadline and thus essentially was lost. In our applications it was not a problem but means of mitigation should be further investigated. The Kafka broker can handle approximately 400,000 spikes per second and this number scales linearly with the number of brokers. There are no limits in principle to the size of a computer cluster and cloud providers put huge clusters at our disposal. We simulated networks with up to 30 neurons for this paper but much larger networks should be possible and will be the focus of further work. In conclusion, we demonstrated key features that make CloudBrain especially suited for some types of experiments and argue that trading small form factor and low power consumption for such extra features can be sensible for research purposes.

## 6 Acknowledgements

We thank Mathias Neerup for invaluable help setting up CloudBrain and for discussions on architecture and implementation. We thank Cao Danh Do and Emil Bonde Larsen for help preparing the robot and its environment. Finally we thank SDU-Biorobotics and The Centre for BioRobotics for funding the project.

